# Transcriptome Analysis of *Aegle marmelos* (L.) Corrêa Fruit and Leaf Tissues: An Underexploited Fruit Tree

**DOI:** 10.1101/2024.04.29.591804

**Authors:** Prashant Kaushik

**Affiliations:** Chaudhary Charan Singh Haryana Agricultural University, Hisar 125004, India

**Keywords:** Bael, *A. marmelos*, denovo assmly, transcriptome

## Abstract

*Aegle marmelos* (L) Correa is both a horticultural and medicinally important tree in South-East Asia. Both of its fruits and leaves have several medicinal values including treatment of certain chronic diseases. Despite its importance, the genomic resource of this tree is very scarce. The objectives of this work were to conduct a de novo transcriptome assembly of leaf tissue from A. *marmelos*, locally known as bael in India. Comparison studies were conducted between the fruit transcriptome and leaf transcriptome of *A. marmelos*. This is the first time that the fruit transcriptome *of A. marmelos* has been reported in the literature. There were 47 million clean reads were generated for fruit tissue and 34 million clean reads of leaf tissue. The reads were assembled to 61, 860 unigenes. Around 83% of fruit transcripts were mapped to the original assembly whereas 89% of leaf transcripts were mapped to the original assembly. Whereas, in terms of molecular function, the gene ontology (GO) terms were related to binding, cellular activity and transporter activity and genes expressed only in leaf and fruit tissue were 14,578 and 11086, respectively, Overall, RNA sequencing may be an effective method for determining the pathway and genes in the underexploited fruit crops.

## Introduction

*Aegle marmelos* (L.) Corrêa, commonly known as Bael, Wood apple, or Stone apple, belongs to the family Rutaceae, an essential native Indian medicinal tree (Orwaetal., 2009) grown near the temple’s gardens. The fruit tree is also found in dry deciduous forests of Sri Lanka, Java, Malaysia, Myanmar and Thailand and is cultivated widely in tropical Africa and southeast Asia (Warrier et al., 2010). Fruits are oval shaped and have a tough rind with orange-coloured pulp. The fruit pulp of the tree has a laxative effect and works as a nostrum for summers. The fruits are used to treat ear problems, diarrhoea, ophthalmia, ulcer, inflammation, stomachalgia, dysentery, cardiopalmus, seminal weakness and respiratory infections (Sharma et al., 2016). The fruit pulp is a rich source of sugars, protein, fiber, fat, minerals, phosphorus, calcium, riboflavin and vitamins C and A (Patil and Muthusamy, 2020).

In agro-processing, fruits of *A. marmelos* are pressed and processed to produce juice, jam, syrup, jelly, and toffee. Apart from nutrients, the fruits are also rich in bioactive compounds like angeline, marmelosin and phenolic compounds. The main phenolic compounds found in the fruit of *A*.*marmelo*s are chlorogenic acid, ellagic acid, etc. (Warrier et al., 2002).

Considering the benefits of *A. marmelos* fruits against several diseases, the selection was made for large-fruited varieties as much as up to 1 kg per fruit. The fruit is of different shape and size like spherical, pyriform, and oval. The fruit size varies between 2 to 8 inches in diameter. The peak fruiting period of *A. marmelos* under Indian conditions is from May to June, with a fruit yield ranging between 50 to 140 kg per tree (Sharma et al., 2007). *Aegle marmelos* fruits are climacteric, and their outer shell turns yellow at maturity. Moreover, based on the variety, the fruits either possess a thin papery or hard woody shell. Overall, varieties with a thin, readily breakable rind, fine-textured sweet pulp, with few seeds are preferred for processing and table purposes. Britto et al. (2009) reported the PCR-based RAPD markers for *A. marmelos* for the first time from Western Ghats of India. As a traditional medicinal plant in Indian and adjoining areas, it is very important to study its genome and transcriptome to focus on the pharmacology and efficacy of its active compounds (Kaushik et.al., 2018).

Therefore, the current study compiled and contrasted the *A. marmelos* leaf and fruit transcriptome to include details on the *A. marmelos* fruit transcriptome in our analysis. For establishing a good and enriched fruit development, it is important to understand the biosynthetic pathways and genomic resources for *A. marmelos*. The information regarding genes and biosynthetic pathways could help establish nutraceutical enriched fruits of *A. marmelos* and differentiate transcriptomes of fruit and leaves of *A. marmelos*.

## Material and Methods

### Plant Material, RNA Extraction, and RNA Sequencing

Leaf sample of *A. marmelos* variety Kagzi was procured from the Government Garden Nursery, Kurukshetra, Haryana, India. Leaf transcriptome data collected for reference from our previous study was downloaded from the NCBI Bioproject (PRJNA433585). At the same time, the fruit samples were collected in mid-April of 2018. Fresh fruits that were commercially ripe were harvested from cultivar “Kagzi”, growing at the Government Garden Nursery (coordinates at 29°58′06.9” N 76°52′50.8” E), Haryana, India. Fruit samples were kept in liquid nitrogen until the RNA was extracted.

### RNA extraction and sequencing

The fruits were opened, and the pulp and fruit flesh were selected for RNA sequencing. The method described by Chang et al. (1993) was considered for isolating total RNA. DNase was used to remove DNA contamination in the RNA samples. Agarose (1%) gel electrophoresis was then performed to get the reliability of the RNA samples. Following that, RNA extracted from these fruits was pooled to create a single sample for the construction of cDNA libraries using the TruSeq RNA Library Prep Kit v2 (Illumina, Inc., USA) (Illumina, Inc., USA). The Agilent 2100 Bioanalyzer (Agilent, USA) and a 1% agarose gel were used to quantify the RNA. The paired-end reads were generated using the HiSeq 2500 (2 x 150 bp chemistry) equipment. To generate mRNA libraries from each sample, 10g of total RNA was used as defined elsewhere (Jiang L et al. 2011). The mRNA library was validated qualitatively and quantitatively using an Agilent Bioanalyzer Chip DNA 1000 series II (Santa Clara, CA, USA) and sequenced using an Illumina HiSeqTM 2000 to generate single-end 100-base pair (bp) sequences. The RNA sequencing data were submitted to the United States of America’s NCBI (National Center for Biotechnology Information) under the BioProject accession number PRJNA433585 (National Center for Biotechnology Information). We assembled these reads using the Trinity assembler version 6-8-2012 with a minimum contig length of 200bp and an optimal k-mer length of 25 for de novo assembly (Grabherr et al. 2011).

### Annotation and classification

BlastX was used to compare all unigenes to the non-redundant sequence (nr) and Swiss-Prot databases, with an e-value cut-off of 1e-5 used to distinguish between the two databases. Analysis of unigenes from the transcriptome of *A. marmelos* was done in conjunction with comparisons to available EST assemblies, leaf coding sequences, and fruit coding sequences, all using a cut-off of 1e-5.

### Results

In this study, we chose two tissues, i.e., fruit pulp and leaves of *A. marmelos*, and the cDNA library was constructed using TruSeq RNA Library Prep Kit. As we are devoid of *A. marmelos*, genome de novo assembly of the transcripts was performed. A total of 49 million and 35 million as raw reads were obtained for leaf and fruit samples of *A. marmelos*. After trimming, the clean reads were 47 million and 34 million for fruit and leaf tissue, respectively (Table 1).

**Table 1.** Reads generated for the leaf and fruit sample of *A. marmelos*.

| Sample | Raw Reads | Clean reads | Error(%) | Q20(%) | Q30(%) | GC(%) |
| --- | --- | --- | --- | --- | --- | --- |
| Fruit | 49,587,436 | 47,718,736 | 0.02 | 96.86 | 92.22 | 46.33 |
| Leaf | 35,489,094 | 34,841,868 | 0.03 | 96.82 | 88.7 | 44.57 |

Also, GC% for all the unigenes was 46.33% for fruit pulp and 44.57% for leaves. There were 86 million nucleotides with a median length 1,077 and an N50 of 2,058 (Table 2).

**Table 2.** Length distribution of transcripts and unigenes.

| Lengths (bp) | Unigenes |
| --- | --- |
| Minimum Length | 201 |
| Mean Length | 1,405 |
| Median Length | 1,077 |
| Maximum Length | 16,700 |
| N 50 | 2,058 |
| N 90 | 661 |
| Total | 86,898,704 |

### Gene Functional Annotation

The information regarding the annotation of genes using different databases was performed to understand annotated and matched genes (Table 3). Out of 61860 unigenes, 71.49% of genes were annotated with the NR database, and 52.20% were annotated with the GO database. Whereas 75.55% were annotated with the NT database, and the most minor 15.68 were annotated with all the databases. The Venn diagram mapped with the database annotation is presented in Figure 2. The areas of matching and similarity between the genes can be easily understood by depicting the Venn diagram (Figure 1).

**Table 3.** The status of annotated genes in the *A. marmelos* denovo transcriptome assembly.

| **Annotation** | **Number of Unigenes** | **Percentage (%)** |
| --- | --- | --- |
| Annotated in NR | 44,227 | 71.49 |
| Annotated in NT | 46,740 | 75.55 |
| Annotated in KO | 17,635 | 28.5 |
| Annotated in SwissProt | 34,371 | 55.56 |
| Annotated in PFAM | 32,152 | 51.97 |
| Annotated in GO | 32,291 | 52.2 |
| Annotated in KOG | 16,722 | 27.03 |
| Annotated in all Databases | 9,702 | 15.68 |
| Annotated in at least one Database | 49,089 | 79.35 |
| Total Unigenes | 61,860 | 100 |

**Figure 1.**
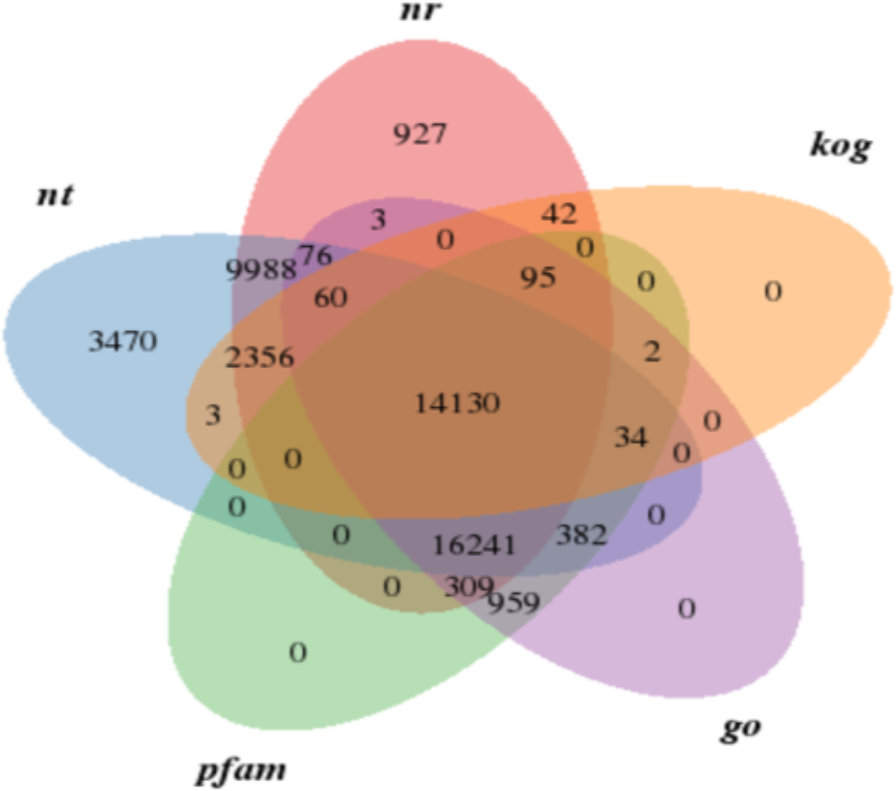
Venn diagram depicting the match between databases in the denovo transcriptome assembly of *A. marmelos*.

**Figure 2.**
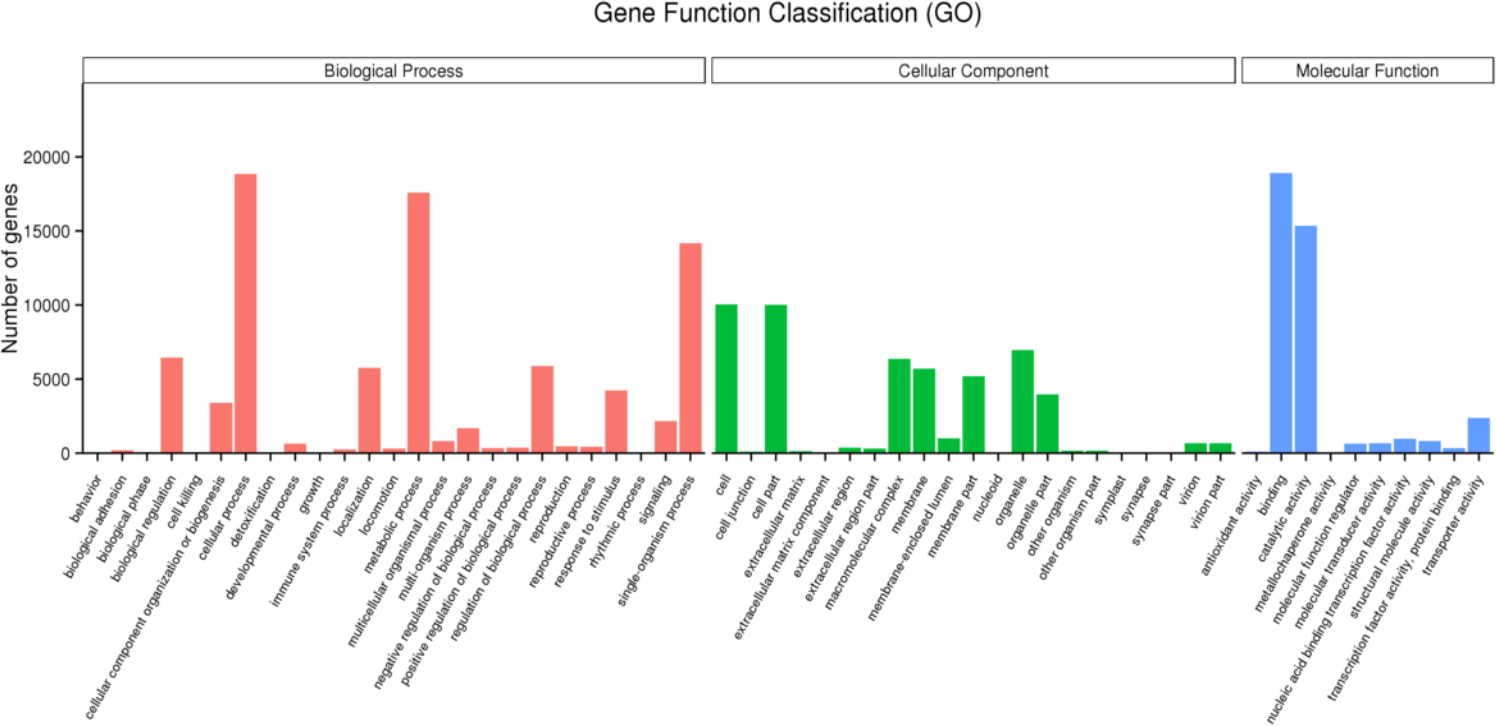
*A. marmelos* transcriptome assembly revealed GO terms for biological mechanism, cellular portion, and molecular function.

### GO and KEGG Classification

Cellular process, metabolic process, and single-organism process were the most expressed GO terms related to biological process (Figure 2). For cellular components, the maximum number of genes were determined for the GO terms cell, cell part and organelle, respectively (Figure 3). Whereas, in terms of molecular function, the GO terms were related to binding, cellular activity and transporter activity (Figure 2). Of all the databases calculated and results obtained from annotation, KEGG classification was used for proper understanding. We can see that different genes are involved in different pathways (Figure 3). The maximum gene expression seen for Translation, was around 1392, lies between 6-8%, whereas the minimum gene expression seen for membrane transport, which was significantly less, i.e., 66, which lies from 0-2% (Figure 3).

**Figure 3.**
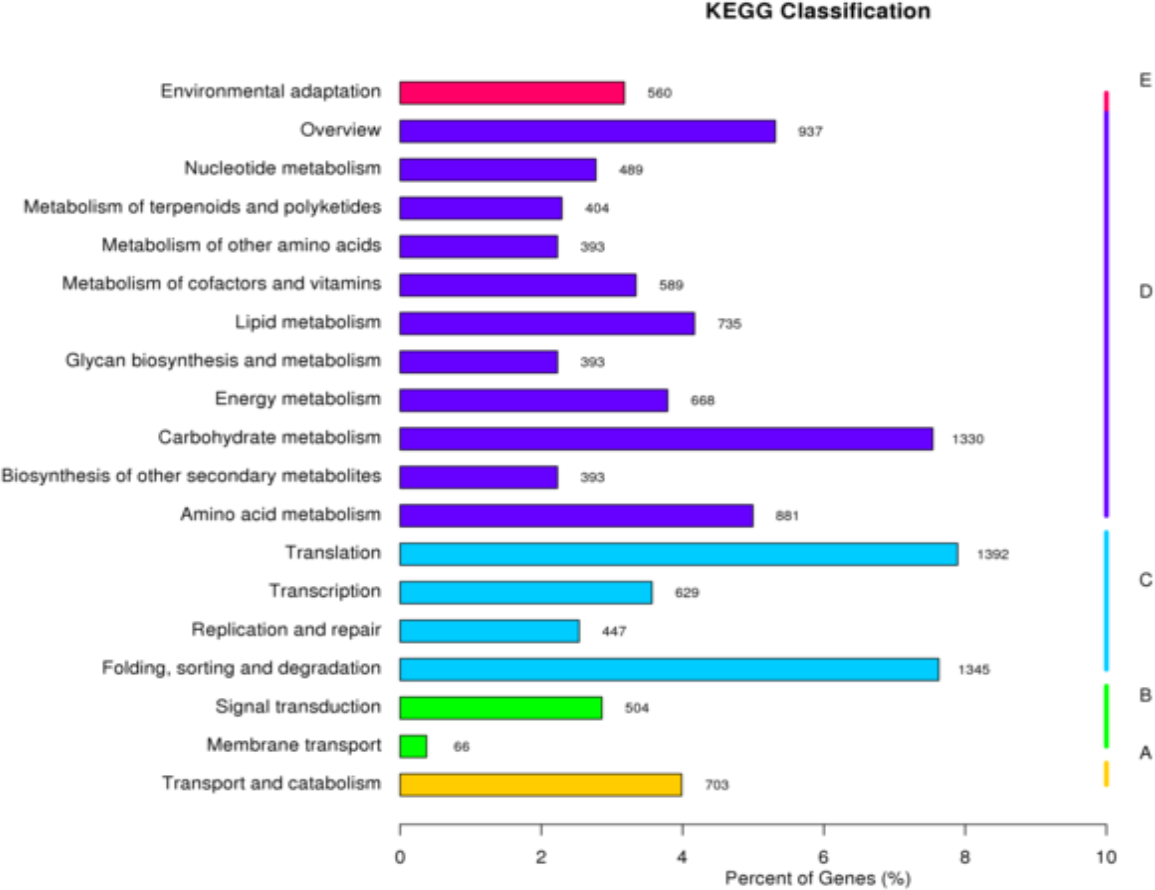
KEGG Classification where Y-axis labels for type of pathway and the X-axis shows the ratio of genes annotated in the pathway versus total genes.

### Expression profiling of unigenes

In order to investigate the differences, reads of leaf and fruit tissue libraries were mapped into the assembly mapped to the special database using the RSEM program. The results revealed that out of a total of 82,560,604 reads generated for leaf and fruit tissue, 39,771,516 (83.35%) of fruits reads were mapped to the database, and 31,090,806 (89.23%) of the leaf reads were mapped to the database (Table 4).

**Table 4.** Reference alignment of the leaf and fruit sample of the *A*.*marmelos*.

| Sample name | Total reads | Total mapped |
| --- | --- | --- |
| Fruit | 47,718,736 | 39,771,516(83.35%) |
| Leaf | 34,841,868 | 31,090,806(89.23%) |

### Filtering the Differential Gene Expression

When there are only two samples or groups, the Venn diagram of expression genes will be plotted. The total number of genes expressed are presented in the Venn diagram. individually in each circle within a group, and the overlap represents the genes expressed in common between groups. Based on the FPKM > 0.3 criteria, the number of genes expressed only in leaf were 14,578 and fruit is 11086, whereas the genes common to both fruit and leaf is around 33,922 (Figure 4).

**Figure 4.**
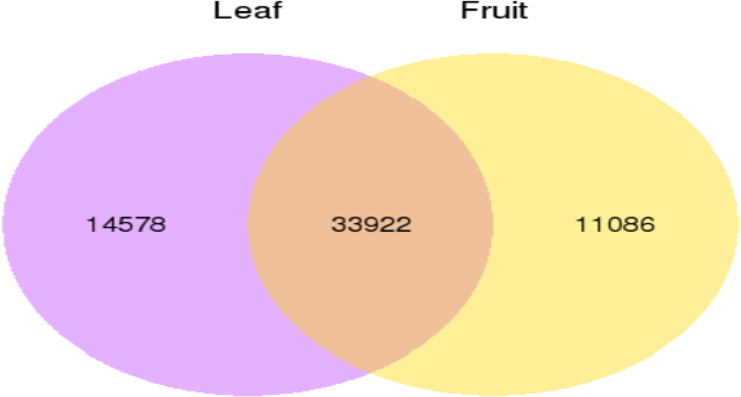
Venn Diagram of expression gene between leaf and fruit tissue of *A. marmelos*.

### KEGG Pathway Enrichment Analysis

The KEG plot revealed the DEG profile enriched in the KEG pathway. The degree of KEG enrichment was determined by Richness Element, q-value, and number of genes (Figure 5). The top20 enriched pathways were laid out in the study (Figure 5). The present study was related to the carbon metabolism, spliceosome, and protein processing in the endoplasmic reticulum (Figure 5). From the KEGG pathway, few mechanisms and functions were determined, wherein the p-value is the same for three mechanisms i.e., for carbon metabolism, pentose phosphate pathway and spliceosome, which is 0.33, whereas the corrected p-value for circadian rhythm plant was 0.079932.

**Figure 5.**
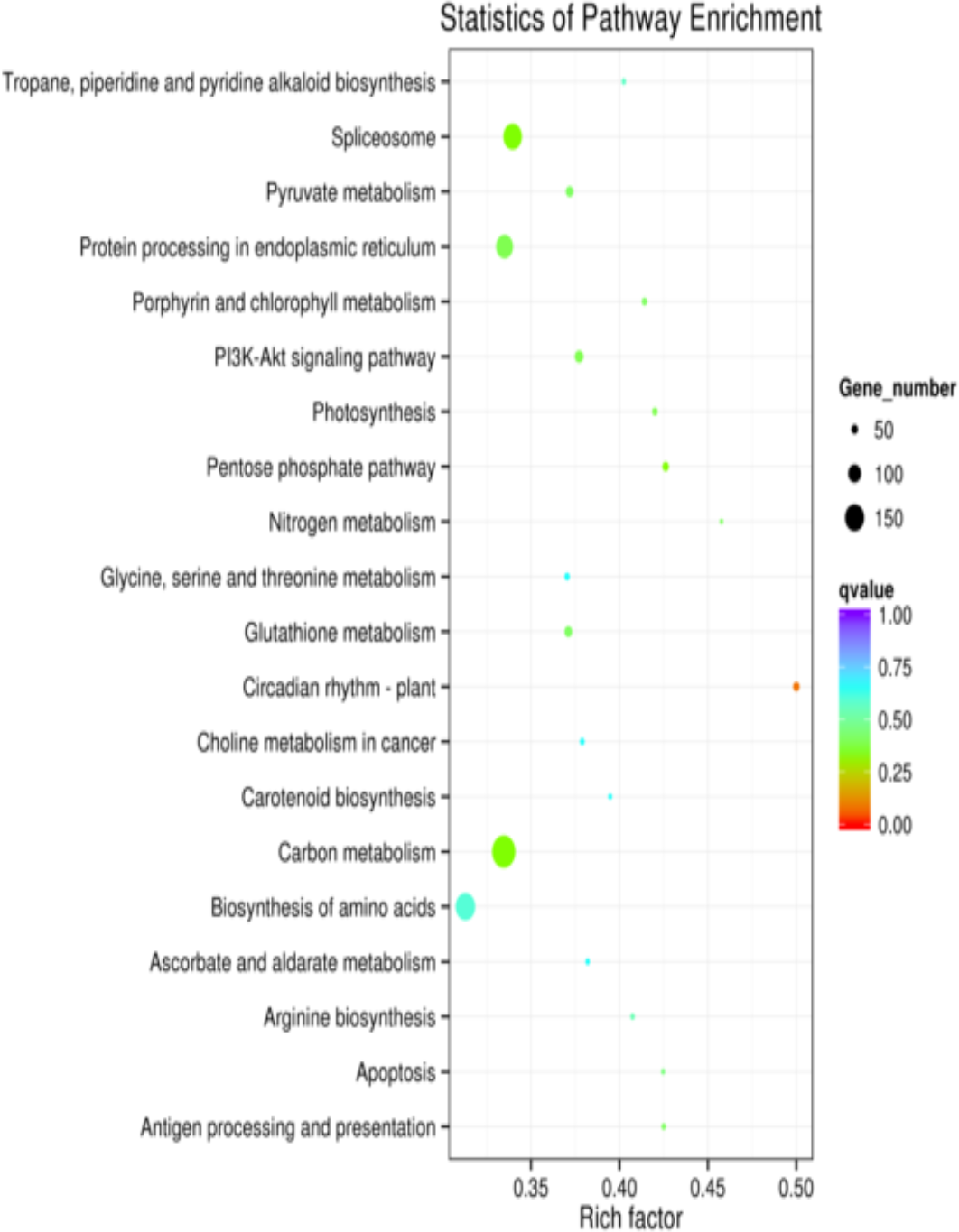
The y axis is the name of the pathway, and the x-axis is the rich. Dot size is the number of the genes, and the colour shows the q value.

## Discussions

In the present study, transcriptome sequencing and de novo assembly of leaf and fruit pulp of *Aegle marmelos* (L.) Corr. Serr (Family: Rutaceae) was performed. Generally, different cultivars show differences in fruit size, weight, yield, sugar content and pulp quantity and seed size, weight, number, and fiber content per fruit (Srivastava and Singh, 2004). That is why its sequencing is essential to know the functioning of several secondary metabolites produced for defence and signalling. With modern tools, several bioactive compounds from almost every plant part of *A. Marmelos* were identified. They obtained transcript sequencing coverage from the plant, fundamental molecular, genetics and biochemical characteristics data can easily be explored. Unigenes from other reported species, such as *Sorbus pohuashanensis*, had 770 bp (Liu et al., 2017). Future transcriptome research may use this information to focus on the genetic pathways that drive fruit development and leaf production of beal, which we expect will be incredibly productive in the future. To discover transcriptionally active genes in bael tissues, which are critical for its production and the quality, this unique transcriptome survey provides a foundation for future in-depth studies on several vital metabolic pathways.

The *A. marmelos* transcriptome’s gene makeup, biological activities, and pathways were studied using 16,722 KOG mapped transcripts. High-throughput sequencing can detect andrographolide production due to many assembled transcripts encoding proteins with over 300 amino acids. The leaf and fruit transcriptome of *A. marmelos* was reported in this study, assisting in identifying several genes involved in secondary metabolite production. The transcripts revealed in this investigation of terpenoid biosynthesis will be helpful in future genetic studies of andrographolide manufacturing routes.

*Myrica rubra* had 531 bp (Feng et al., 2012), and *Platycladus orientalis* had 534 bp (Chang et al., 2012) are inferior to those of our studied unigenes. These sequences are like the unigenes annotated and classified against the NCBI NR database. Therefore, choosing the leaf and fruit pulp for comparative transcriptome analysis will significantly facilitate dissection of the genes involved in organ-specific secondary metabolite biosynthesis. Microsatellite markers revealed 100% polymorphism in genotypes of *A. marmelos* (Sharma and Sharma, 2015; Sivalingam et al., 2016; Kaushik and Kumar, 2018). Based on transcriptome sequencing, we annotated CYP450s (Cytochrome P450 monooxygenases) and Glycosyltransferase (GT). CYP450s and GT constitute the largest and complex superfamily, useful in terpenoid biosynthesis, play vital roles in producing secondary metabolites and various natural products (Weitzel et al., 2015; Rasool et al., 2016; Qi et al., 2017). The leaves and seeds have many other constituents *viz*., limonene, ethanol, dichloromethane, β-phellandrene, methanol, p-cymene, petroleum ether, arabinose, n-hexane, oleic acid, β-caryophyllene, linolenic acid, oleic palmitic acid, cryptone and Humulene which work against several groups of bacteria (Jamal *et al*., 2017; Mahomoodally *et al*., 2018). Dutta *et al*., 2014 extracted aegeline, lupeol, citronella, skimmianine, marmesinine, eugenol, cineol, and cuminaldeyde (from leaves); marmine, and fagarine (from bark); tannin, marmelosin, psoralen, aurapten, luvangetin, and marmelide (from fruit).

Studies have revealed that the Bael tree has many Coumarins like marmenol, marmelosin, scoparone, imperatorin, methyl ether, marmin, xanthotoxol, marmesin, scopoletin, marmelid, psoralen, and umbelliferone (Korkina and Afanasev, 1997; Farooq, 2005). Besides this, several alkaloids like fragrine, aegelenine, aeglin, dictamine, etc., are found in Bael (Manandhar et al., 1978). Screening suitable genotypes for biotic and abiotic stress resistance is needed to identify existing elite genotypes that need to be exploited for further genetic improvement. Bael fruit is rich in phytochemicals essential, and hence several experiments are conducted to understand the leaf and fruit benefits of Bael fruit so that it can be beneficial to humankind. Several changes in morphology and genetic characters are to develop high yielding and highly nutritious fruits.

## Conclusion

Along with the high nutritional value of its fruits, there are several medicinal uses in Ayurveda, such as the cure of jaundice, cancer, and heart alignments. Nevertheless, the *A. marmelos* is not well studied on the lines of molecular biology and genomics. But, still, there is no information about the transcriptome of fruit mRNA. Therefore, in our study, we have tried to assemble and compare the leaf and fruit transcriptome of *A. marmelos*. For the first time, we have compared the fruit transcriptome with the leaf transcriptome of *A. marmelos* where the fruit transcriptome was inferior to the leaf. This work has developed necessary genomic resources for *A. marmelos*. The critical information regarding genes and biosynthetic pathways will help to establish nutraceutical enriched fruits.

